# Progressively reduced cerebral oxygen metabolism and elevated plasma NfL levels in the zQ175DN mouse model of Huntington’s disease

**DOI:** 10.1101/2025.03.16.643551

**Authors:** Qian Wu, Minmin Yao, Hongshuai Liu, Aaron Kakazu, Yuxiao Ouyang, Chang Liu, Ruoxuan Li, Fan Yang, Anqi Wang, Sharmane Surasinghe, Damien Gerochi, Barbara Baldo, Stefanie Jahn, Haiying Tang, Hanzhang Lu, Zhiliang Wei, Wenzhen Duan

**Affiliations:** Division of Neurobiology, Department of Psychiatry and Behavioral Sciences, Johns Hopkins University School of medicine, Baltimore, Maryland, USA; Evotec SE, Manfred Eigen Campus, Hamburg, Germany; CHDI Management for CHDI Foundation, Princeton, NJ, USA; The Russell H. Morgan Department of Radiology and Radiological Sciences, Johns Hopkins University School of Medicine, Baltimore, Maryland, USA; Solomon H. Snyder Department of Neuroscience, Johns Hopkins University School of Medicine, Baltimore, Maryland, USA; Program in Cellular and Molecular Medicine, Johns Hopkins University School of Medicine, Baltimore, Maryland, USA

**Author notes:** These authors contribute to the work equally (Qian Wu, Minmin Yao). **Correspondence to**: Wenzhen Duan, Division of Neurobiology, Department of Psychiatry and Behavioral Sciences, Johns Hopkins University School of Medicine. CMSC 5120, 600 North Wolfe Street, Baltimore, MD 21287.

**Keywords:** Huntington’s disease, mouse model, cerebral oxygen metabolism, brain atrophy;mi neurofilament light chain

## Abstract

Huntington’s disease (HD) is a progressive neurodegenerative disorder caused by a CAG-repeat expansion in exon-1 of the *huntingtin* gene. Currently, no disease-modifying therapies are available, with a significant challenge in evaluating therapeutic efficacy before clinical symptoms emerge. This highlights the need for early biomarkers and intervention strategies. Therefore, it is essential to develop and characterize accurate mouse models and identify early biomarkers for preclinical therapeutic development. In this study, we characterized the pathological progression of the heterozygous zQ175 neo-deleted knock in (zQ175DN) mouse model across four age groups: 3, 6, 10, and 16 months to identify human translatable outcome measures. T2-relaxation-under-spin-tagging (TRUST) MRI was used to assess global CMRO_2_, while T2-weighted MRI was used to analyze brain volumes. Significant brain volume loss was detected as early as 6 months of age, worsening progressively with age in the zQ175 DN mice, resembling HD brain volumetric changes. A decline in CMRO_2_ was observed in 6-month-old zQ175 DN mice, with significant and progressive reductions in 10- and 16-months old HD mice. Additionally, PHP1-positive mutant huntingtin (mHTT) aggregates were present in the striatum of zQ175 DN mice at all four age groups, with intranuclear localization prior to 6 months, transitioning to both intranuclear and neuropil aggregates in older zQ175 DN mice, suggesting that the localization of mHTT aggregates may reflect the severity of HD pathogenesis. Interestingly, plasma neurofilament light chain (NfL) protein concentrations were significantly elevated at 6 months of age and older zQ175DN mice. These findings provide valuable insights for selecting outcome measures in preclinical evaluations of HD therapies using the zQ175 DN mouse model.

## Introduction

Huntington’s disease (HD) is a progressive neurodegenerative disorder caused by a CAG repeat expansion in exon 1 of the Huntingtin (*HTT*) gene, leading to motor, cognitive, and psychiatric abnormalities. Medium spiny neurons in the striatum are particularly vulnerable to mutant huntingtin HTT protein (mHTT), resulting in striatal atrophy before clinical symptoms appear. A key pathological hallmark of HD brain is the aggregation of mHTT, first observed in HD mouse models and then confirmed in HD patients’ brains. Although the clinical diagnosis of HD is based on motor symptoms, brain functional changes and volumetric loss can occur decades prior to clinical symptoms^1–6^. MRI, being noninvasive, has played pivotal role in uncovering brain alterations and facilitating the identification of biomarkers across disease progression in HD^7^.

HD mouse models have been instrumental in understanding disease pathogenesis^8^ and offer valuable tools for developing therapeutics and biomarkers. The zQ175DN heterozygous mouse model is a refined knock-in mouse model, where exon 1 of human *HTT* gene (with ∼190 CAG repeats) is integrated into the murine *Htt* homolog gene. Unlike the zQ175 model, the zQ175DN model has undergone removal of the floxed neo-cassettes to eliminate potential artificial effects on the symptomatic and pathological development in the HD model. Previous studies suggest that this deletion increases mutant HTT expression^9,10^, while reducing phenotypic variability, therby improving experimental reproducibility^11^. Consequently, the zQ175DN mouse model offers additional advantages, including that removal of the neo cassette simplifies the genetic background, reduces confounding effects, and enhances the consistency of experimental outcomes by minimizing genetic variability.

The evaluation of therapeutic efficacy requires sensitive measures that can be used repeatedly over time, particularly when clinical symptoms have not yet appeared. Volumetric MRI detection of brain volume loss appears up to two decades before clinical motor diagnosis^12–14^. Additionally, metabolic disturbances are increasingly recognized in HD pathogenesis^15–20^. Recent single cell RNA-sequencing studies have shown impaired mitochondrial metabolism in the most vulnerable striatal neurons before motor symptoms manifest in mouse models of HD, with similar changes in HD human brains^21^. Brain imaging studies using MRI, MR spectroscopy, and PET/SPECT have demonstrated that metabolic changes occur in the premanifest HD^1–4,6,22^. PET studies have indicated that these metabolic changes precede and progress more rapidly than brain volume changes in premanifest HD^22^. MRS studies also highlight significant metabolic changes in the HD brain^23–28^. As a result, monitoring metabolic changes using noninvasive measures might serve as a biomarker for tracking disease progression and evaluating therapeutic efficacy in premanifest HD. Moreover, premanifest HD patients exhibit reduced cerebral metabolic rate of oxygen (CMRO_2_) in response to visual stimulation, supporting early-stage metabolic dysfunction in HD^29^.

In this study, we conducted a comprehensive characterization of the heterozygous zQ175 DN model using human-translatable MRI measures to analyze CMRO_2_ and brain volumetric changes across four age groups, 3, 6, 10, and 16 months, representing different stages of disease progression. Additionally, mHTT aggregates were evaluated using PHP-1 antibody^30^ and plasma NfL levels were measured post-MRI scans in the same cohorts. Our results indicated that, similar as HD patients, brain atrophy appears to be a stable biomarker in the zQ175DN model. The significantly reduced CMRO_2_ was evident at 10 months and older HD mice. The decline trend was observed at 6 months old HD mice, but there was not statistically different due to larger variability in this measure. mHTT aggregates progressively increased, expanding from the intranuclear location to cell bodies and neuropils. Interestingly, plasma NfL levels were significantly elevated in 6 months and older zQ175 DN mice. These findings suggest that zQ175 DN is a valuable HD tool for preclinical studies and provide guidance for selecting appropriate outcome measures to assess therapeutic efficacy in this model.

## Materials and Methods

### Mice and experimental procedures

The mouse experimental protocols involved in this study were approved by the Johns Hopkins University Animal Care and Use Committee and were conducted in accordance with the National Institutes of Health guidelines for the care and use of laboratory animals. Male and female heterozygous zQ175 DN KI (JAX stock #: 029928) and age-matched wild-type (WT) controls were obtained from the Jackson Laboratory. The heterozygous mice carry one wild-type mouse *HTT* allele and a CAG-expanded human *HTT* exon 1 sequence with 190 ± 21 trinucleotide repeats. The zQ175DN KI mice used in the experiments have been backcrossed with C57BL/6J inbred mice (JAX stock #: 000664) as the background strain. All mice were housed at room temperature (23 degrees Celsius, 30-70% humidity) in a 12-hour light-dark reversed cycle, specific pathogen-free facility with free access to food and water. Tail biopsies were obtained from all mice and sent to Laragen Inc (Culver City, CA, now a part of Transnetyx Inc) to confirm genotypes and CAG size.

*In vivo* MRI scans were conducted in the cohorts of mice at the following ages, 3, 6, 10, and 16 months. All MRI measures were performed in the mouse’s dark phase. Following the completion of MRI scans, mice were anesthetized using isoflurane (VetOne, Fluriso), transcardial perfusion was performed with chilled phosphate-buffered saline (PBS; Corning, 46-013-CM) and then 4% paraformaldehyde (PFA; Sigma-Aldrich, 158127) in PBS for histological studies. After careful extraction of brain from the animals’ skulls, the perfused brains were placed into 4% PFA overnight at 4 °C to allow additional fixation, then cryoprotected using 30% sucrose (Sigma-Aldrich, S0389) in 0.02% sodium azide-PBS (Sigma-Aldrich, S2002). The fixed brains were positioned in tissue freezing medium (Leica, 14020108926), sliced into 30 μm coronal sections using a cryostat (Leica, CM1950), and stored free-floating in 0.02% sodium azide-PBS for immunohistology examination.

### T2-relaxation-under-spin-tagging (TRUST) MRI

MRI scans were conducted using a horizontal-bore 11.7 T Bruker BioSpec instrument (Bruker, Ettlingen, Germany) equipped with an actively shielded field gradient with a maximum intensity of 0.74 T/m. A 72-mm quadrature volume resonator was the transmitter, and a four-element (2 x 2) phased-array coil was the receiver. Mice were initially anesthetized with 2% vaporized isoflurane mixed in medical air. After induction, the mouse was transferred to a motorized, water-heated animal bed, where its head was positioned within the coil element using a bite bar and secured with two ear pins. Anesthesia induction was at 1.5% concentration for 15 minutes, followed by maintenance at 1.0%. The respiration rate of each mouse was continuously monitored during experiments to ensure survival and maintain consistent respiratory rates across all experimental mice. If a mouse exhibited a breathing rate exceeding 150 breaths per minute, the maintenance isoflurane dose was slightly increased to 1.2%. This anesthesia protocol has been previously utilized and documented ^31^. Additionally, each mouse was immobilized using a bite bar and ear pins, then placed on a water-heated animal bed with temperature control.

A subject-specific field map was acquired to homogenize the B0 field, followed by a global, second-order shimming. Following a localizer scan to determine the optimal imaging slice and viewing parameters, a multi-slice rapid acquisition with relaxation enhancement (RARE) sequence was used to obtain T2-weighted anatomical images of the whole brain: repetition time (TR)/time to echo (TE) = 3000/11 ms, field of view (FOV) = 15 mm × 15 mm, matrix size = 128 × 128, spatial resolution = 0.1 mm × 0.1 mm, slice thickness = 0.5 mm, RARE factor = 4, 35 coronal slices, and scan duration = 1.6 min.

To determine the oxygen extraction fraction (OEF), we utilized a previously reported TRUST protocol^32^. Briefly, an axial time-of-flight (TOF) angiogram sequence was used to locate the mouse’s arterial and venous blood vessels: TR/TE = 20/2.7 ms, FOV = 15 mm × 15 mm, matrix size = 256 × 256, 25 axial slices, slice thickness = 0.5 mm, and scan duration = 2.0 min. The optimal location of imaging slice was centered around the sinus confluence, which was empirically determined by a previous study to offer the best signal-to-noise ratio. Then, the TRUST scan was conducted using the following parameters: TR/TE = 3500/6.5 ms, FOV = 16 mm × 16 mm, matrix size = 128 × 128, voxel size = 125 μm × 125 μm, slice thickness = 0.5 mm, inversion slab thickness = 2.5 mm, post-labeling delay (PLD) = 1000 ms, effective time-to-echo (eTE) = 0.25, 20, 40 ms, and scan duration = 2.8 min.

The cerebral blood flow was observed with phase-contrast (PC) MRI that was detailed in a previous study^33^. In short, a coronal TOF angiogram was performed to locate the left/right internal carotid arteries and basal artery, the major blood suppliers into the brain. The following parameters were used: TR/TE = 45/2.6 ms, FOV = 25 mm × 16 mm, matrix size = 256 × 256, slice thickness = 0.5 mm, 11 coronal slices, and scan duration = 3.2 min. Then, three sagittal TOFs were completed, each positioned along one of the arteries: TR/TE = 60/2.5 ms, FOV = 16 mm × 16 mm, matrix size = 256 × 256, slice thickness = 0.5 mm, and scan duration per artery = 20 sec. These angiograms were aligned with the coronal TOF to position imaging slices perpendicular to each vessel’s blood flow. Using the reported encoding velocity (VENC) of 20 cm/s ^29^, the blood flow velocity was determined via PC with the following parameters: TR/TE = 15/3.2 ms, FOV = 15 mm × 15 mm, matrix size = 300 × 300, slice thickness = 0.5 mm, receiver bandwidth = 100 kHz, partial Fourier acquisition factor = 0.7, flip angle = 25°, and scan duration per artery = 0.6 min.

### TRUST MRI image analysis

TRUST MRI scan data was collected using animal IDs and analyzed by investigators blinded to age groups and genotypes. Using a custom-written graphic-user-interface (GUI) tool on MATLAB (MathWorks, Natick, MA) from previous studies^32,33^, the OEF was processed. Briefly, the program facilitated pairwise subtractions between the control and labeled images. Investigators manually drew a region of interest (ROI) around the sinus confluence on the difference image. The four voxels with the most significant difference signals were spatially averaged within the ROI. The signal intensities from the difference images at three different eTEs (0.25, 20, and 40 ms) were fitted into a mono-exponential curve to determine the blood T2 value. The T2 was then converted to the venous blood oxygenation, Yv, using a previously described T2-Yv calibration plot^34^. The OEF is defined as the difference in oxygen concentration between the arterial and venous blood, encapsulated by the following relationship^35^:

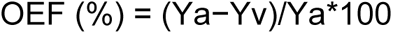

Arterial blood oxygenation tends to be close to saturation^36^, so we assumed a value of 0.99 for Y_a_, the arterial oxygen saturation fraction, to calculate the OEF.

Similarly, cerebral blood flow (CBF) was calculated using custom-written scripts on MATLAB (MathWorks, Natick, MA) and methodology from a previous study^33^. Using the complex-difference image from PC scans on the left/right internal carotid and basilar arteries, investigators manually drew ROIs around each artery of interest. The grayscale arterial voxels in the ROI were converted to blood flow (ml/min) using a pre-determined phase velocity map. The total blood flow (TBF) to the brain was the sum of the blood flow in these three feeding arteries. The unit-volume CBF, normalized to the subject-specific brain volume, was calculated by dividing the TBF by the product of the brain volume and brain density (1.04 g/ml)^37^.

Using the CBF and venous oxygenation obtained from TRUST MRI, we can estimate the CMRO_2_ based on a prior study^3^. Briefly, the relationship between CMRO_2_ and CBF was described using the Fick principle:

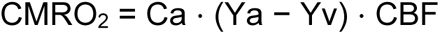

C_a_ refers to the molar concentration of oxygen in a unit volume of blood and was assumed to be 882.1 μmol O_2_/100 ml blood based on the literature^38,39^.

### Structural MRI image analysis

The analysis of the structural MRI images closely followed the methodology detailed in our previous studies^40,41^. Briefly, processing of MRI images began by adjusting the scan orientation as defined by the Paxinos mouse brain atlas to rigidly align the orientation of the template image using automated image registration (AIR; http://www.bmap.ucla.edu/portfolio/software/AIR/) and DiffeoMap software (https://www.mristudio.org/). Here, the isotropic resolution of our subject image was also adjusted to 0.1 mm × 0.1 mm × 0.1 mm per pixel to match the template image. The template image used in this study was a single image from an age-matched littermate control mouse with the median brain volume within the control group. The skull-stripping feature in ROIEditor software (https://www.mristudio.org/) removed non-brain tissue signals. Investigators validated this automated processing by manually checking each image slice and removing any non-brain tissue signals as necessary with careful reference to the mouse brain atlas.

Using DiffeoMap, the skull-stripped subject images had their signal intensity values of the gray/white matter and cerebral spinal fluid normalized to those of the template image using an affine model, piece-wise linear function. Following this, the intensity histograms of the subject and template images were matched to be as similar as possible. DiffeoMap sent the intensity-normalized, rigidly aligned scan images to a Linux cluster that conducted Large Deformation Diffeomorphic Metric Mapping (LDDMM). LDDMM relies on an internal mouse brain atlas consisting of in vivo population-averaged T2-weighted mouse brain images with carefully delineated major gray and white matter structures^28^. The values of the LDDMM transformations were calculated using ROIEditor, yielding a quantitative measurement that can be converted to brain volumes in mm^3^.

### Immunofluorescent staining and quantification of mHTT aggregates

The brain sections were washed and permeabilized 3 times for 10 minutes each in TBST (0.5% Triton X-100 (Sigma-Alrich, T9284) in tris-buffered saline (Corning, 46-012-CM). Then, brain sections were incubated with a blocking buffer consisting of 5% normal goat serum (Abcam, ab7481) in TBST for 1 hour at room temperature. Afterward, the sections were incubated in PHP1 antibody (Mouse anti-HTT-Q20, clone 27A7, CH01950, kindly provided by CHDI contracted Coriell Institute for Medical Research, 1:500 dilution) with blocking buffer overnight at 4 °C. After three 10-minute washes in TBST, the sections were incubated in the dark with a working dilution of Alexa 555-conjugated secondary antibody (Goat anti-mouse, 1:1000, ThermoFisher, A-21422) in blocking buffer for 1 hour at room temperature. After thoroughly washing with TBST, the sections were counterstained in a 5 µg/ml dilution of Hoechst 33342 (Sigma-Aldrich, B2261) for 15 minutes. Immunofluorescence images were acquired using a Zeiss Axio Observer an inverted microscope and LSM700 confocal microscope.

To quantify the extent of mHTT aggregates, images were obtained using a 20× objective from each brain section’s left and right cortices, striata, and hippocampi in triplicate, six images per brain region. Six mouse brains/genotype. The representative images of each group were obtained using a 63× objective from LSM700 confocal microscope. All images were analyzed using Fiji ImageJ software, with careful control for consistent thresholding to determine mHTT-positive counts.

### Neurofilament light chain (NfL) quantification

The plasma samples were collected before the perfusion and kept at -80°C and thawed on assay day on ice. The Simoa NF-LIGHT v2 Advantage Kit (Lot # 504139) was applied to determine NfL levels in plasma. NfL quantitation was performed in technical duplicates, at a dilution of 1:20. On the day of the assay the samples were thawed on ice and centrifuged for 5 min at 10000 × g at 4 °C. The standard calibrators were provided in the advantage v2 kit (Lot# 504139) at a final concentration range between 4.44 and 281 pg/mL. Each run included 3 quality control (QC) samples to assess the technical performance of the assay: Control 1 (Low QC, 4.44 pg/mL) and Control 2 (High QC, 281 pg/mL) were provided within the kit; an additional control (Middle QC, 28.14 pg/mL) was generated as a 1:10 dilution of Control 2 in sample diluent. For the final analysis, recovery of standard and QC samples was calculated as percentage of their back calculated concentration over their nominal concentration.

Acceptance criteria were considered when the back-calculated data of at least 2 out of three controls would fall within 25% variation of the expected values. All controls passed the acceptance criteria.

Similarly to the QCs, the duplicate measurements of the samples were back calculated to the respective calibrator curve to determine the pg/mL NfL concentrations and adjusted according to the dilution factor.

### Statistical Analysis

Data in the figures are represented as the mean ± standard deviation (SD) unless otherwise noted. Investigators were blinded to gender, genotype, and age groups when processing and analyzing results. Research personnel were rigorously trained and supervised to standardize ROI selection criteria in determining the structural MRI brain volumes, OEF, CBF, and CMRO_2_. Statistical analyses were performed using Prism software (version 10.2.3; GraphPad Software, Boston, MA). For the quantification of mHTT, a two-tailed, unpaired Student’s t-test was used in comparing different age of mHTT aggregation, and a two-tailed, paired Student’s t-test was used in comparing the difference of intranuclei and extranuclei mHTT at different ages. Two-tailed unpaired Student’s *t*-tests were performed to analyze statistical significance in structural MRI brain volumes, OEF, CBF, and CMRO_2_ between WT and HD groups.

The NfL data were analyzed by the Simoa software using a 4-parameter logistic curve fit, 1/y² weighted. Each data set was transformed by log10 and tested for normal distribution (Gaussian distribution: D’Agostino-Pearson). The groups were then compared using the Mixed-effects model (REML) test followed by post-hoc Tukeýs multiple comparisons test. P-values less than 0.05 were considered as statistically significant, with *p<0.05, **p<0.01, ***p<0.001, and ****p<0.0001.

## Data Availability

All datasets associated with this study are included in the paper. Any queries for raw data, in-house data processing scripts, and other original materials should be made to. These requests will be promptly reviewed to determine whether the application is subject to intellectual property or confidentiality requirements. Any data and materials that can be shared will be released after executing a material transfer agreement.

## Results

### Age, gender, and CAG size in zQ175 DN mice

In this study, we examined zQ175DN mice at four distinct ages, 3, 6, 10, and 16 months, for cross-sectional characterization. Equal numbers of male and female mice were included in each genotype and age groups. Individual data points are shown, and blue color represent male data and red color represent female data in all figures. These age groups were chosen to capture stages before, during, and after the onset of HD-like phenotypes. The CAG repeat size in the experimental mice was 190 ± 21.

### Progressive brain atrophy in zQ175 DN mice

To investigate whether HD-like progressive brain atrophy occurs in zQ175DN mice, structural MRI (sMRI) was performed on mice at 3, 6, 10, and 16 months of age. Automatic segmentation of brain volumes was conducted, with quantification performed using LDDMM as described in the methods. Representative sMRI images from zQ175DN and WT mice at the indicated ages, along with segmented brain regions for volumetric analysis, are shown in Figure 1A.

**Figure 1.**
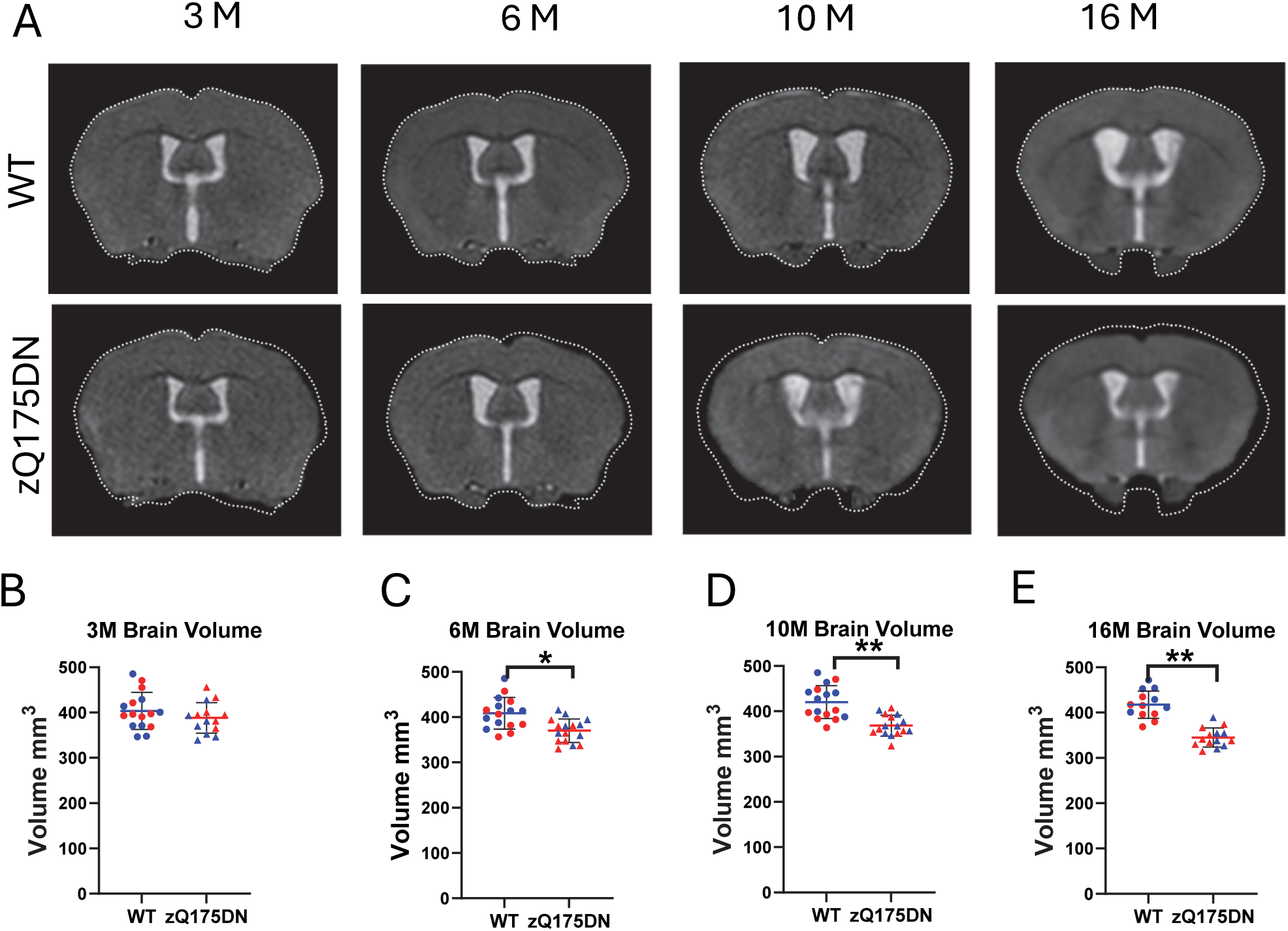
Progressive brain atrophy in zQ175DN mice. (**A)** Representative MRI images in mouse brains at indicated genotypes and ages. (**B-E**) Whole brain volume in the zQ175DN mice and age-matched WT controls at 3, 6, 10, and 16 months. Blue symbols represent male data, and red symbols indicate female data. Individual data points are shown. *p < 0.05; **p < 0.01 by Student’s *t*-test.

At 3 months, no significant differences were observed in the volumes of the neocortex (Figure 1B), though zQ175DN mice showed trends towards smaller volumes. Significant atrophy was detected in the whole brain (Figure 1C) in zQ175DN mice at 6 months. The atrophy progressed with age, becoming more pronounced at 10 (Figure 1D) and 16 months (Figures 1E). These findings confirm that sMRI measures of brain volume are sensitive and reliable indicators of brain atrophy in zQ175DN HD mice. No gender-dependent differences were noticed in brain volumes of zQ175DN model.

### Reduced cerebral oxygen metabolism in zQ175 DN mice

TRUST MRI was performed following procedures outlined in the methods to assess global CMRO_2_ in zQ175DN mice at 3, 6, 10, and 16 months, along with age- and sex-matched controls. OEF and CBF were calculated, and unit-mass CMRO_2_ was derived from OEF and CBF using the Fick principle. At 3 months, no significant differences in CMRO_2_ or OEF were observed between zQ175DN and WT mice (Figures 2A, E). However, a trend toward reduced CMRO_2_ and OEF appeared in zQ175DN mice by 6 months (Figures 2B, F). Significant reductions in both parameters were seen at 10 and 16 months (Figures 2C, D, G, H). Notably, global CBF remained stable across all ages in zQ175DN mice (Figures 2I-L). None of these measures show gender-dependent differences.Therefore, we conducted statistics by combining data from both genders. Our data suggest that CMRO_2_ could be a useful noninvasive biomarker for disease progression in zQ175DN HD mice, as the same MRI technique has been established in human brain^42^.

**Figure 2.**
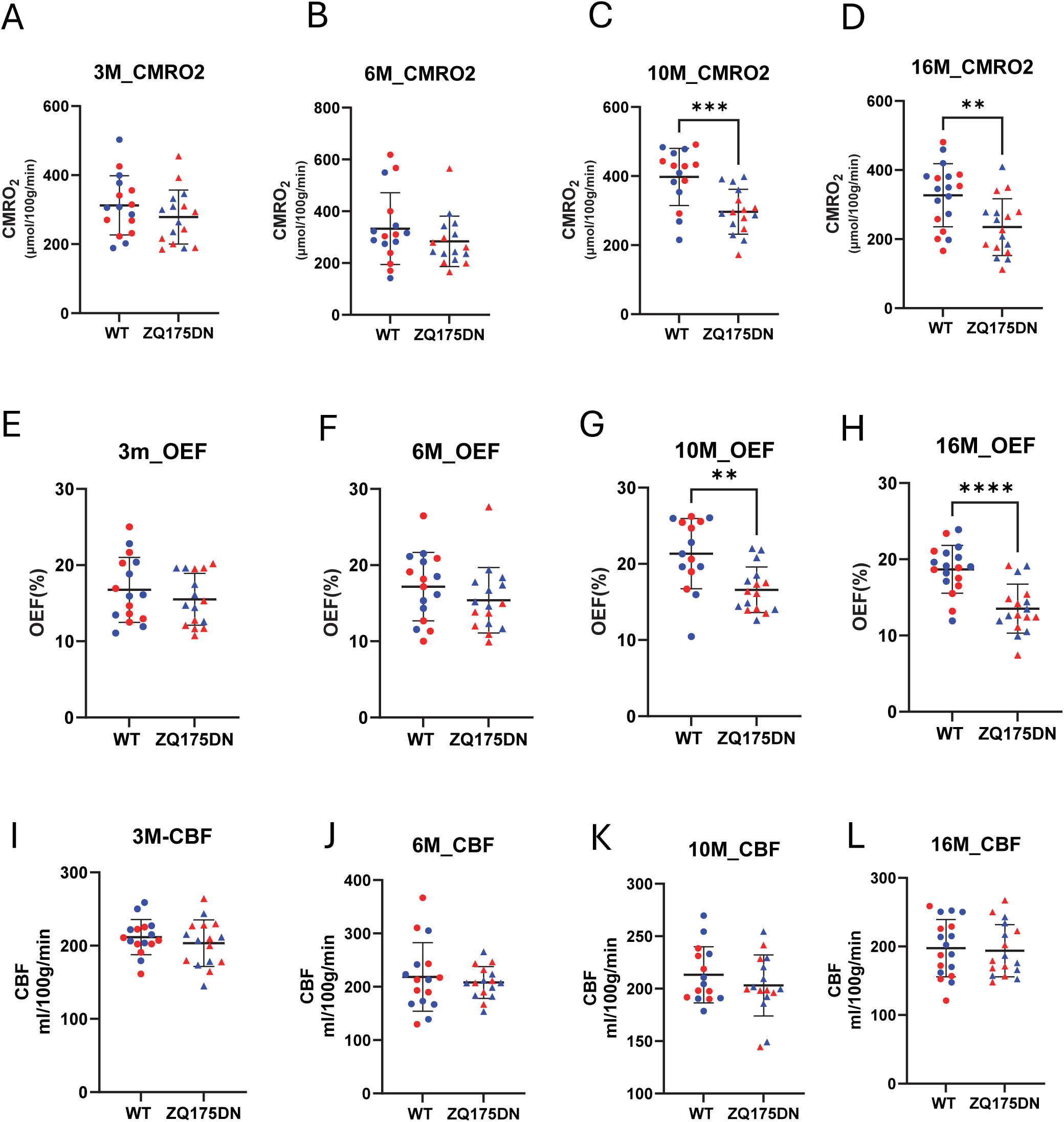
Altered cerebral metabolic rate of oxygen (CMRO_2_) in zQ175DN mice. **(A-D)** Global CMRO_2_ values in zQ175DN mice and age-matched wild type (WT) controls at 3, 6, 10, and 16 months (M). (**E-H**) Oxygen extraction fraction (OEF) in the zQ175DN mice and age-matched WT controls at 3, 6, 10, and 16 months (M). (**I-L**) global cerebral blood flow (CBF) in the zQ175DN mice and age-matched WT controls at 3, 6, 10, and 16 months (M). Blue symbols represent male data, and red symbols indicate female data. Individual data points are shown. **p < 0.01; ***p < 0.001, ****p<0.0001 by Standard Student’s *t*-test.

### PHP1 positive mHTT aggregates are evident in the brain of zQ175 DN mice

Pathogenic exon 1 of the mHTT protein consists of three domains: the N-terminal region, the polyQ sequence, and the C-terminal proline-rich domain (PRD)^43^. The mHTT protein can form soluble monomers, oligomers, and fibril-rich insoluble inclusion bodies (IB) that accumulate in both the cytoplasm and nucleus ^44^. mHTT aggregation begins with primary nucleation, where spontaneous aggregation nuclei induce conformational changes in other proteins, leading to their incorporation into the aggregate through templated aggregation^44^. Expansion of the polyQ region in mHTT accelerates nucleation and aggregation, recruiting additional monomers to form fibrils^45^. Fibrillary branching occurs specifically in mHTT aggregates, unlike wild-type HTT, which does not naturally form fibrillary structures^43^. The PRD of mHTT is the most dynamic region in assembled fibrils, promoting misfolding and the evolution of new conformations^46,47^. The C-terminal polyproline repeat facilitates the assembly of unbound fibrils^48^.

In our study, we used the PHP1 antibody, which targets the polyproline region of exon 1 in HTT fibrils and recognizes mHTT aggregates. PHP1-positive mHTT aggregates were detected in several brain regions, particularly in the striatum of all four age groups of zQ175DN mice. Initially, these aggregates were mainly localized to the nucleus, with the striatum being the most affected region (Figure 3A). As the disease progressed from 6 to 10 months, the number of PHP1-positive aggregates increased (Figure 3B), the localization of mHTT aggregates are expanding from the nucleus to cytoplasma and neuropils (Figure 3C). Interestingly, no further increase in PHP1-positive aggregates was observed in 16-month-old zQ175DN mice (Figure 3B-C), while extranuclei mHTT aggregates were more than intranuclei mHTT in 16-month-old zQ175DN mice (Figure 3C).

**Figure 3.**
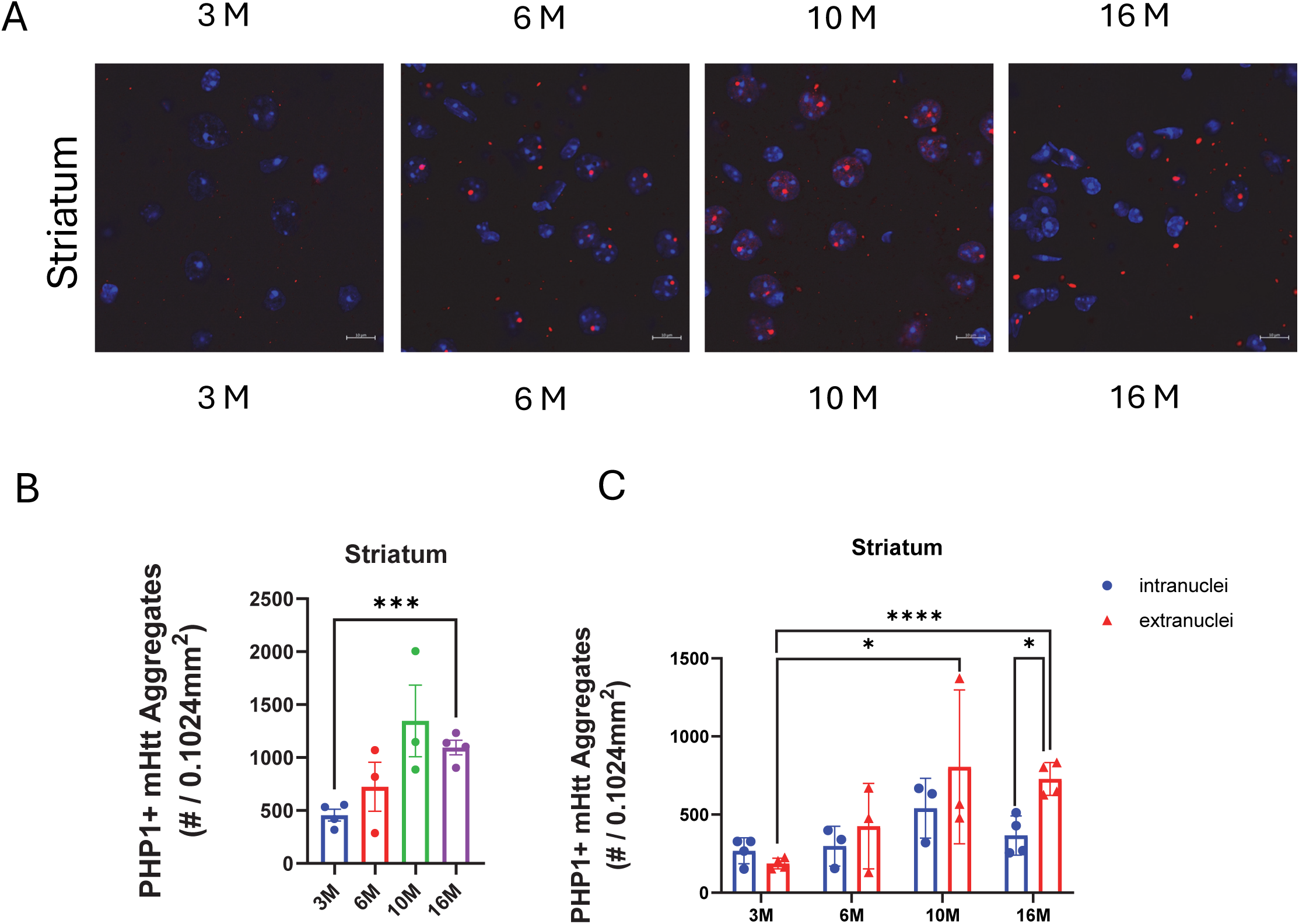
PHP1-positive mHTT aggregates in zQ175DN mice. **(A**) Representative images of mHTT aggregates (red) detected by PHP1 antibody in the striatum of zQ175DN mice at 3, 6, 10 and 16 months (M) of age. Hoechst labels nucleus (blue). Scale bar = 10 µm. (**B**) Quantification of all PHP1 positive mHTT aggregates in the striatum of zQ175DN at 3, 6, 10 and 16 months of age. (**C**) Quantification of intranuclear and extranuclear mHTT aggregates in the striatum of zQ175DN mice at 3, 6, 10 and 16 months of age. Individual data points are shown. *p<0.05, ***p < 0.001, ****p<0.0001 by Standard Student’s *t*-test.

### Plasma Neurofilament light chain (NfL) is increased in zQ175 DN mice

NfL has emerged as a significant prognostic biomarker in HD, offering insights into neuronal damage and disease progression^49^. Its elevated levels in bodily fluids indicate neuronal injury. In HD, measuring NfL levels provides valuable information about disease onset, progression, and potential therapeutic interventions^50,51^. NfL levels were measured in the plasma of zQ175DN mice and their littermate controls across four age groups. A rising trend in plasma NfL levels was observed in zQ175DN mice aged 6 months and older, though high variability was noted in the NfL measurements from these samples (Figure 4A). After Log10 transformation, the data passed the normality test (D’Agostino & Pearson test) and revealed statistically significant increases in plasma NfL levels in zQ175DN mice at 6, 10, and 16 months of age (Figure 4B).

**Figure 4.**
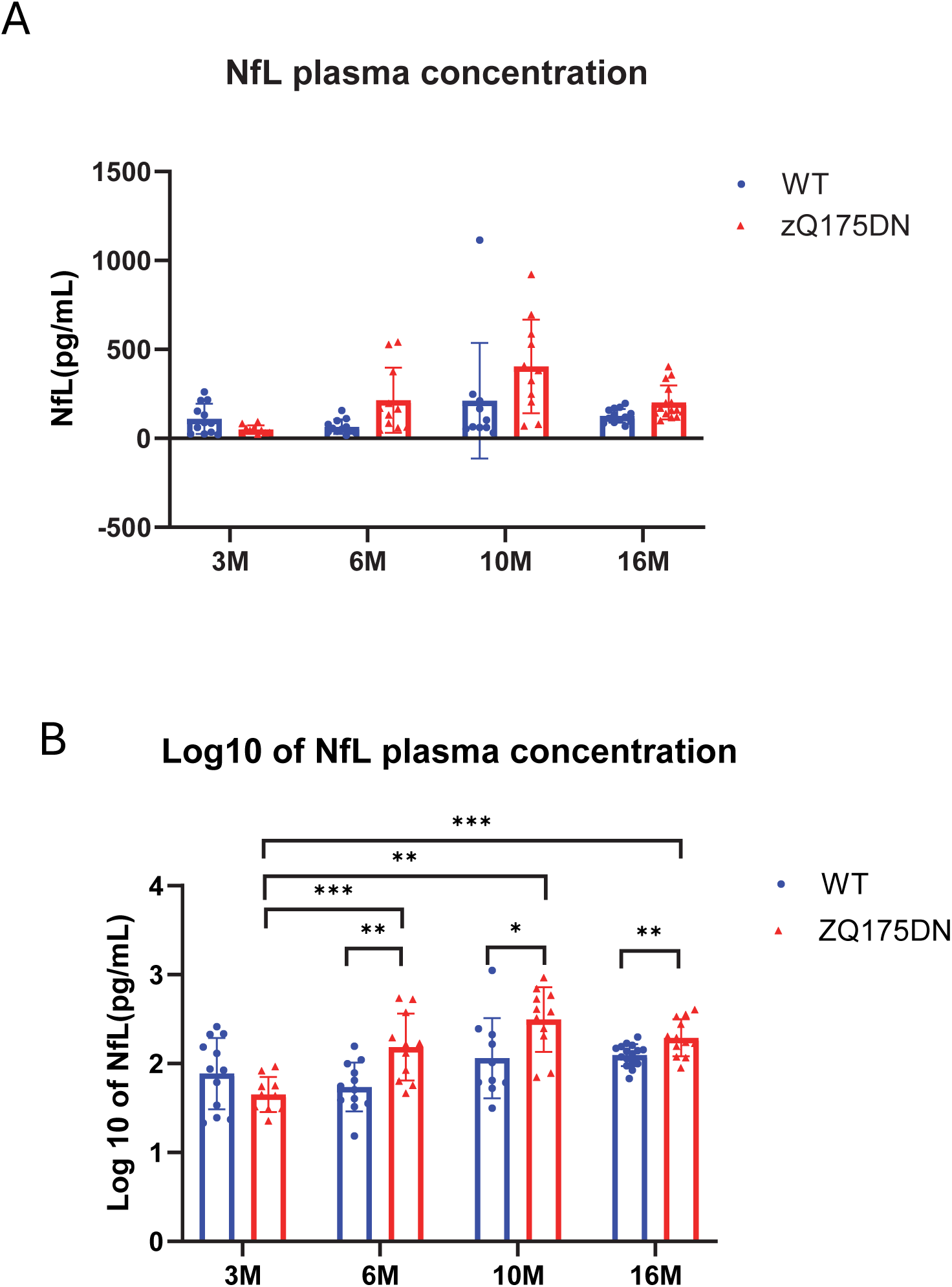
Plasma NfL levels are increased in zQ175 DN mice. **(A**) Quantification of NfL levels in mouse plasma from zQ175DN mice (red) and age-matched controls (blue). Data are shown as Mean with SD. (**B**) Log10 transformation of NfL values in mouse plasma from zQ175DN mice (red) and age-matched controls (blue. Each data set was transformed by log10 and tested for normal distribution (Gaussian distribution: D’Agostino-Pearson). The groups were then compared using the Mixed-effects model (REML) test followed by post-hoc Tukeýs multiple comparisons test. *p<0.05, **p<0.01, ***p<0.001.

## Discussion

Our study highlights the utility of the zQ175DN mouse model in replicating the progressive nature of HD across multiple parameters, positioning it as a valuable mouse model for preclinical evaluation of therapeutic interventions. We confirmed progressive brain atrophy in zQ175DN mice. These observations align with the degenerative patterns reported in previous studies using the zQ175 model, further validating the zQ175DN model for HD research.

Importantly, the study considered the impact of sex on disease progression. Our data showed no significant sex-dependent differences in brain volume changes in this model. Brain atrophy consistency across both male and female zQ175DN mice strengthens the reliability of this model for generalizing findings across sexes. These results underscore the model’s robustness for assessing HD-related brain atrophy and its potential for testing therapeutic approaches aimed at slowing or halting disease progression.

Although our data showed a downward trend in CMRO_2_ as early as 6 months and even 3 months of age, these changes did not reach statistical significance. A significant decline in CMRO_2_ in zQ175DN mice was observed at 10 months and later age, which occurs later than the onset of detectable brain atrophy. This delayed reduction in CMRO_2_ suggests a dissociation between structural and functional declines in HD progression. One explanation for the delayed changes in cerebral oxygen metabolism than volumes loss is the inherent variability in metabolic measures, which tends to be higher compared to brain volume changes. This variability indicates that a larger sample size might be required to improve the statistical power and reliability of the functional assessments in mice. We found no sex-dependent differences in CMRO_2_ values in zQ175DN mice, further supporting the use both male and female mice in this model.

The accumulation of mHTT aggregates within the nucleus and neuropil regions of the striatum underscores the pathological relevance of the zQ175DN model in HD research. Early-stage zQ175DN mice exhibit primarily intranuclear aggregates, a pattern that shifts with disease progression as neuropil aggregates become more prominent in older mice. This transition from localized nuclear inclusion to widespread neuropil involvement likely reflects the worsening severity of HD, with increasing cellular dysfunction as the disease advances. These observations suggest that intranuclear mHTT aggregates may represent an early hallmark of HD pathology, while neuropil aggregates signify a more advanced state of neuronal damage and synaptic dysfunction. The zQ175DN model thus provides a valuable framework for investigating the dynamics of mHTT aggregation and the cellular mechanisms underlying HD progression.

NfL is an established biomarker for neurodegeneration, reflecting axonal damage and neuronal injury. The analysis revealed a progressive elevation in plasma NfL levels in zQ175DN mice beginning at 6 months of age. This pattern aligns with disease progression, with statistically significant elevations detected at 6, 10, and 16 months of age, suggesting ongoing neuronal damage as the disease advances. The rising trend in plasma NfL observed here corresponds with the emergence of HD-like brain atrophy in zQ175DN mice, supporting its utility as a biomarker for monitoring neuronal injury/ neurodegeneration in HD. Although NfL levels were elevated, the high variability in measurements, particularly in later stages, indicates the need for further refinement in sampling or control for confounding variables. Log10 transformation improved the normality of the data, enhancing the reliability of statistical interpretations. These findings emphasize the importance of NfL as a prognostic marker for neurodegenerative changes and highlight the zQ175DN mouse model is valuable for tracking disease progression and therapeutic efficacy.

It is important to highlight that both structural and cerebral oxygen metabolic MRI measures employed in this study are fully adaptable for use in the human brain, underscoring the translational potential of findings from this mouse model. The non-invasive nature of MRI allows for continuous monitoring changes in the brain, providing a robust tool for assessing HD progression. This capability enhances the potential of MRI measures to be seamlessly integrated into HD clinical trials, effectively bridging the gap between preclinical findings and clinical applications. Our study demonstrates the utility of the zQ175DN HD mouse model in capturing the complex dynamics of HD-related neurodegeneration, making it a valuable platform for the preclinical evaluation of therapeutic interventions.

Moreover, by offering a comprehensive assessment of both structural alterations and metabolic dysfunctions, as well as mHTT aggregates and plasma NfL levels, the current study strengthens the zQ175DN model’s relevance in preclinical research. These findings indicate the importance in using multiple biomarkers, which could significantly improve the precision of diagnosis of disease onset, the monitoring of disease progression, and the assessment of treatment efficacy in human HD patients.

## Acknowledgements

This project is supported by CHDI Foundation Inc, NIH R01 NS127344, NIH R01 NS124084 (to W.D). R01AG081932 (to Z.W), R01AG071515 (to H.L).

## Author contributions

QW conducted sMRI and immunohistochemistry experiments and analyzed results. MY conducted TRUST MRI experiments and analyzed results. MY, HLiu, AK, YO, CL, RL, FY, AW, SS, DG, HLu, ZW contributed to MRI scan sequence development, MRI scans and data analysis, brain tissue and plasma sample collections. HT contributed to overall experimental design, mouse age selection, and manuscript discussion. BB and SJ conducted NfL assay and data analysis. WD conceptualized the study. WD, QW, MY wrote the manuscript.

## References

1. Andrews TC, Weeks RA, Turjanski N, et al. Huntington’s disease progression. PET and clinical observations. Brain. 1999;122 ( Pt 12):2353–2363.

2. Ciarmiello A, Cannella M, Lastoria S, et al. Brain white-matter volume loss and glucose hypometabolism precede the clinical symptoms of Huntington’s disease. J Nucl Med. 2006;47(2):215–222.

3. Feigin A, Tang C, Ma Y, et al. Thalamic metabolism and symptom onset in preclinical Huntington’s disease. Brain. 2007;130(Pt 11):2858–2867.

4. Lopez-Mora DA, Camacho V, Perez-Perez J, et al. Striatal hypometabolism in premanifest and manifest Huntington’s disease patients. Eur J Nucl Med Mol Imaging. 2016;43(12):2183–2189.

5. Paulsen JS. Functional imaging in Huntington’s disease. Exp Neurol. 2009;216(2):272–277.

6. Young AB, Penney JB, Starosta-Rubinstein S, et al. PET scan investigations of Huntington’s disease: cerebral metabolic correlates of neurological features and functional decline. Ann Neurol. 1986;20(3):296–303.

7. Vasilkovska T, Verschuuren M, Pustina D, et al. Evolution of aberrant brain-wide spatiotemporal dynamics of resting-state networks in a Huntington’s disease mouse model. Clin Transl Med. 2024;14(10):e70055.

8. Menalled L, Brunner D. Animal models of Huntington’s disease for translation to the clinic: best practices. Mov Disord. 2014;29(11):1375–1390.

9. Southwell AL, Smith-Dijak A, Kay C, et al. An enhanced Q175 knock-in mouse model of Huntington disease with higher mutant huntingtin levels and accelerated disease phenotypes. Hum Mol Genet. 2016;25(17):3654–3675.

10. Wu J, Mohle L, Bruning T, et al. A Novel Huntington’s Disease Assessment Platform to Support Future Drug Discovery and Development. Int J Mol Sci. 2022;23(23).

11. Speziale R, Montesano C, Di Pietro G, et al. The Urine Metabolome of R6/2 and zQ175DN Huntington’s Disease Mouse Models. Metabolites. 2023;13(8).

12. Bates GP, Dorsey R, Gusella JF, et al. Huntington disease. Nat Rev Dis Primers. 2015;1:15005.

13. Ross CA, Aylward EH, Wild EJ, et al. Huntington disease: natural history, biomarkers and prospects for therapeutics. Nat Rev Neurol. 2014;10(4):204–216.

14. Scahill RI, Andre R, Tabrizi SJ, Aylward EH. Structural imaging in premanifest and manifest Huntington disease. Handb Clin Neurol. 2017;144:247–261.

15. Browne SE. Mitochondria and Huntington’s disease pathogenesis: insight from genetic and chemical models. Ann N Y Acad Sci. 2008;1147:358–382.

16. Su B, Wang X, Zheng L, Perry G, Smith MA, Zhu X. Abnormal mitochondrial dynamics and neurodegenerative diseases. Biochim Biophys Acta. 2010;1802(1):135–142.

17. Quintanilla RA, Johnson GV. Role of mitochondrial dysfunction in the pathogenesis of Huntington’s disease. Brain Res Bull. 2009;80(4-5):242–247.

18. Li XJ, Orr AL, Li S. Impaired mitochondrial trafficking in Huntington’s disease. Biochim Biophys Acta. 2010;1802(1):62–65.

19. Reddy PH, Shirendeb UP. Mutant huntingtin, abnormal mitochondrial dynamics, defective axonal transport of mitochondria, and selective synaptic degeneration in Huntington’s disease. Biochim Biophys Acta. 2012;1822(2):101–110.

20. Mochel F, Haller RG. Energy deficit in Huntington disease: why it matters. J Clin Invest. 2011;121(2):493–499.

21. Lee H, Fenster RJ, Pineda SS, et al. Cell Type-Specific Transcriptomics Reveals that Mutant Huntingtin Leads to Mitochondrial RNA Release and Neuronal Innate Immune Activation. Neuron. 2020;107(5):891–908 e898.

22. Tang CC, Feigin A, Ma Y, et al. Metabolic network as a progression biomarker of premanifest Huntington’s disease. J Clin Invest. 2013;123(9):4076–4088.

23. Mochel F, N’Guyen TM, Deelchand D, et al. Abnormal response to cortical activation in early stages of Huntington disease. Mov Disord. 2012;27(7):907–910.

24. van den Bogaard SJ, Dumas EM, Teeuwisse WM, et al. Exploratory 7-Tesla magnetic resonance spectroscopy in Huntington’s disease provides in vivo evidence for impaired energy metabolism. J Neurol. 2011;258(12):2230–2239.

25. Unschuld PG, Edden RA, Carass A, et al. Brain metabolite alterations and cognitive dysfunction in early Huntington’s disease. Mov Disord. 2012;27(7):895–902.

26. Zacharoff L, Tkac I, Song Q, et al. Cortical metabolites as biomarkers in the R6/2 model of Huntington’s disease. J Cereb Blood Flow Metab. 2012;32(3):502–514.

27. Tkac I, Henry PG, Zacharoff L, et al. Homeostatic adaptations in brain energy metabolism in mouse models of Huntington disease. J Cereb Blood Flow Metab. 2012;32(11):1977–1988.

28. Jenkins BG, Koroshetz WJ, Beal MF, Rosen BR. Evidence for impairment of energy metabolism in vivo in Huntington’s disease using localized 1H NMR spectroscopy. Neurology. 1993;43(12):2689–2695.

29. Klinkmueller P, Kronenbuerger M, Miao X, et al. Impaired response of cerebral oxygen metabolism to visual stimulation in Huntington’s disease. J Cereb Blood Flow Metab. 2021;41(5):1119–1130.

30. Ko J, Isas JM, Sabbaugh A, et al. Identification of distinct conformations associated with monomers and fibril assemblies of mutant huntingtin. Hum Mol Genet. 2018;27(13):2330–2343.

31. Wei Z, Xu J, Chen L, et al. Brain metabolism in tau and amyloid mouse models of Alzheimer’s disease: An MRI study. NMR Biomed. 2021;34(9):e4568.

32. Wei Z, Xu J, Liu P, et al. Quantitative assessment of cerebral venous blood T(2) in mouse at 11.7T: Implementation, optimization, and age effect. Magn Reson Med. 2018;80(2):521–528.

33. Wei Z, Chen L, Lin Z, et al. Optimization of phase-contrast MRI for the estimation of global cerebral blood flow of mice at 11.7T. Magn Reson Med. 2019;81(4):2566–2575.

34. Li W, van Zijl PCM. Quantitative theory for the transverse relaxation time of blood water. NMR Biomed. 2020;33(5):e4207.

35. O’Brien C, Okell TW, Chiew M, Jezzard P. Volume-localized measurement of oxygen extraction fraction in the brain using MRI. Magn Reson Med. 2019;82(4):1412–1423.

36. Lin AL, Qin Q, Zhao X, Duong TQ. Blood longitudinal (T1) and transverse (T2) relaxation time constants at 11.7 Tesla. MAGMA. 2012;25(3):245–249.

37. Bothe HW, Bodsch W, Hossmann KA. Relationship between specific gravity, water content, and serum protein extravasation in various types of vasogenic brain edema. Acta Neuropathol. 1984;64(1):37–42.

38. Chong SP, Merkle CW, Leahy C, Srinivasan VJ. Cerebral metabolic rate of oxygen (CMRO2) assessed by combined Doppler and spectroscopic OCT. Biomed Opt Express. 2015;6(10):3941–3951.

39. Ulatowski JA, Oja JM, Suarez JI, Kauppinen RA, Traystman RJ, van Zijl PC. In vivo determination of absolute cerebral blood volume using hemoglobin as a natural contrast agent: an MRI study using altered arterial carbon dioxide tension. J Cereb Blood Flow Metab. 1999;19(7):809–817.

40. Liu H, Zhang C, Xu J, et al. Huntingtin silencing delays onset and slows progression of Huntington’s disease: a biomarker study. Brain. 2021;144(10):3101–3113.

41. Peng Q, Wu B, Jiang M, et al. Characterization of Behavioral, Neuropathological, Brain Metabolic and Key Molecular Changes in zQ175 Knock-In Mouse Model of Huntington’s Disease. PLoS One. 2016;11(2):e0148839.

42. Xu F, Ge Y, Lu H. Noninvasive quantification of whole-brain cerebral metabolic rate of oxygen (CMRO2) by MRI. Magn Reson Med. 2009;62(1):141–148.

43. Jarosinska OD, Rudiger SGD. Molecular Strategies to Target Protein Aggregation in Huntington’s Disease. Front Mol Biosci. 2021;8:769184.

44. Wagner AS, Politi AZ, Ast A, et al. Self-assembly of Mutant Huntingtin Exon-1 Fragments into Large Complex Fibrillar Structures Involves Nucleated Branching. J Mol Biol. 2018;430(12):1725–1744.

45. Boatz JC, Piretra T, Lasorsa A, Matlahov I, Conway JF, van der Wel PCA. Protofilament Structure and Supramolecular Polymorphism of Aggregated Mutant Huntingtin Exon 1. J Mol Biol. 2020;432(16):4722–4744.

46. Isas JM, Langen R, Siemer AB. Solid-State Nuclear Magnetic Resonance on the Static and Dynamic Domains of Huntingtin Exon-1 Fibrils. Biochemistry. 2015;54(25):3942–3949.

47. Lin HK, Boatz JC, Krabbendam IE, et al. Fibril polymorphism affects immobilized non-amyloid flanking domains of huntingtin exon1 rather than its polyglutamine core. Nat Commun. 2017;8:15462.

48. Shen K, Calamini B, Fauerbach JA, et al. Control of the structural landscape and neuronal proteotoxicity of mutant Huntingtin by domains flanking the polyQ tract. Elife. 2016;5.

49. Voysey ZJ, Owen NE, Holbrook JA, et al. A 14-year longitudinal study of neurofilament light chain dynamics in premanifest and transitional Huntington’s disease. J Neurol. 2024;271(12):7572–7582.

50. Rodrigues FB, Byrne LM, Tortelli R, et al. Mutant huntingtin and neurofilament light have distinct longitudinal dynamics in Huntington’s disease. Sci Transl Med. 2020;12(574).

51. Byrne LM, Rodrigues FB, Blennow K, et al. Neurofilament light protein in blood as a potential biomarker of neurodegeneration in Huntington’s disease: a retrospective cohort analysis. Lancet Neurol. 2017;16(8):601–609.

